# Single-cell mass distributions reveal simple rules for achieving steady-state growth

**DOI:** 10.1101/2023.02.02.526759

**Authors:** Benjamin R.K. Roller, Cathrine Hellerschmied, Yanqi Wu, Teemu P. Miettinen, Scott R. Manalis, Martin F. Polz

## Abstract

Optical density is a common method for measuring exponential growth in bacterial batch cultures. However, there is a misconception that such exponential growth is equivalent to steady-state growth, which is a distinct physiological state that improves experimental reproducibility. Determining precisely when steady-state growth occurs is technically challenging and is aided by paired single-cell and population-level measurements. Using microfluidic mass sensors and optical density, we explore when in typical laboratory batch cultures steady-state growth occurs. We show that cell mass increases by an order of magnitude within a few hours of dilution into fresh medium and that steady-state growth is only achieved when cultures are inoculated with high dilutions from overnight stationary phase cultures. At high dilutions, *Escherichia coli* and *Vibrio cyclitrophicus* grown in different rich media achieve steady-state growth approximately 4 total biomass doublings after inoculation. We can decompose these dynamics into 3 doublings of average cell mass and 1 doubling of cell number for both species. We also show that batch cultures in rich media depart steady-state growth early in their growth curves at low cell and biomass concentrations. Achieving and maintaining steady-state growth in batch culture is a delicate balancing act, and we provide general guidance for commonly used rich media. Quantifying single-cell mass outside of steady-state growth is an important first step towards understanding how microbes grow in their natural context, where fluctuations pervade at the scale of individual cells.

**Importance:** Microbiologists have watched clear liquid turn cloudy for over 100 years. While the cloudiness of a culture is proportional to its total biomass, growth rates using such optical density measurements are challenging to interpret when cells change size. Many bacteria adjust their size at different steady-state growth rates, but also when shifting between starvation and growth. Optical density cannot disentangle how mass is distributed among cells of different sizes, and directly measuring how mass is distributed among cells has been a major challenge. Here we use single-cell mass measurements to demonstrate that a population of cells in batch culture achieves a stable mass distribution for only a short period of time. Achieving steady-state growth in rich medium requires low initial biomass concentrations and enough time for the coordination of individual cell and population growth. Steady-state growth is important for reliable cell mass distributions in a culture and we discuss how mass variation outside of steady-state can impact physiology, ecology, and evolution experiments.

---

Tracking microbial growth via optical density (OD) is fundamental and simple(1, 2), but it can also mislead by obscuring the processes underlying growth. A culture exhibiting exponential OD growth is often falsely equated with steady-state growth, despite ample literature stating otherwise(2–5). Bacterial cells alter their mass, external dimensions, and macromolecular composition as the nutrient availability and chemistry of the growth medium change in batch culture(4–8). The average cell only exhibits a consistent macromolecular composition when the population achieves balanced and steady-state growth. Steady-state growth describes when the frequency distributions of all measurable cellular properties are time-invariant and it implies balanced growth, which is when all constituent parts of a cell are synthesized at the same exponential rate(3). OD is a proxy for the total mass concentration in a liquid medium(2, 9) and does not distinguish between changes in the cell number, average cell mass, or mass variation among cells. Therefore, OD measurements alone obscure key physiological transitions in batch culture, such as the entry into steady-state growth.

Verifying steady-state or balanced growth is laborious, but essential for examining bacterial cell size regulation(10, 11), cell cycle progression(12–14), and cell organization(15). Many independent aspects of growth must be measured for each culture while repeatedly performing dilutions to guarantee nutrients are available in excess of biosynthetic demand. In practice, many studies simply allow for a certain number of OD doublings (typically at least 10) in unchanging conditions to ensure cells are in steady-state(2, 16). Steady-state growth can also be assessed by measuring the mass distribution of individual cells in a growing population. If the frequency distribution of cell mass in a growing population is time-invariant then steady-state has effectively been achieved(2, 3). Observing a constant average mass is also a good approximation of steady-state growth, though exceptions can exist if transient changes to growth, division, and death serendipitously balanced one another out. Ensuring steady-state growth is undoubtedly important, but less attention has been given to quantifying the coordination of cell mass and cell number dynamics during the transition into steady-state.

Here we ask if simple guidelines can ensure bacterial cultures are reliably in steady-state growth and quantify how cells change as they transition into and out of steady-state growth. We measured three aspects of growth in batch culture: single-cell mass distributions, cell concentration, and optical density. We used microfluidic mass sensors to measure single-cell buoyant mass(17, 18) and cell concentration simultaneously. Buoyant mass is the product of a cell’s volume and the density difference between cell material and its surrounding fluid. Changes in cellular buoyant mass can be directly translated to changes in cellular dry mass using dry density, which is the density of a cell’s dry material(17). We use mass in place of buoyant mass for clarity throughout the rest of this manuscript, unless specifically noted otherwise. We grew two types of bacteria in commonly used rich, undefined growth media: a typical laboratory bacterium (*Escherichia coli* K12 MG1655 in Lysogeny Broth at 37°C) and a recently isolated marine bacterium which has been minimally passaged in the laboratory (*Vibrio cyclitrophicus* 1G07 in Marine Broth 2216 at 25°C). This allows us to examine if there are general rules for achieving and departing from steady-state growth among diverse bacteria in commonly used cultivation conditions.

The ability of a culture to achieve steady-state growth depends on the initial OD, that is mass concentration, in the culture. We performed dilutions of 1:100, 1:1,000, or 1:10,000 from overnight stationary phase cultures into fresh medium to vary the initial OD, then measured population dynamics over about 5 hours. The mean cell mass immediately increased in all cultures with no apparent lag time upon encountering fresh medium (Figure 1 a-f, Figure 2 a-f), while cell number did not increase for about 2.5 hours (Figure 1 g-i, Figure 2 g-i). The immediate mean cell mass increase is also reflected as rapid changes in OD (Figure 1i, Figure 2i). *V. cyclitrophicus* and *E. coli* achieved steady-state growth in the 1:10,000 dilution cultures because they exhibited 3 properties over a sustained period: time-invariant mass distributions, a constant mean cell mass, and near-parallel increases in OD and cell number (steady-state highlighted by gray boxes in Figure 1 and 2). While *V. cyclitrophicus* briefly achieved steady-state in the 1:1,000 dilution cultures (Figure 1 b, e, h), *E. coli* did not (Figure 2 b, e, h), and neither species achieved steady-state in the 1:100 dilution cultures (Figure 1 c, f, i, Figure 2 c, f, i) despite having obvious exponential increases in OD (Figure 1i, Figure 2i). The rate of nutrient depletion in a 1:100 dilution culture is presumably too rapid for the coordination of single-cell and population growth, because mean cell mass decreases before the first population doubling occurs (Figure 1f & i, Figure 2f & i). Large dilutions are necessary to reliably achieve steady-state growth.

**Figure 1:**
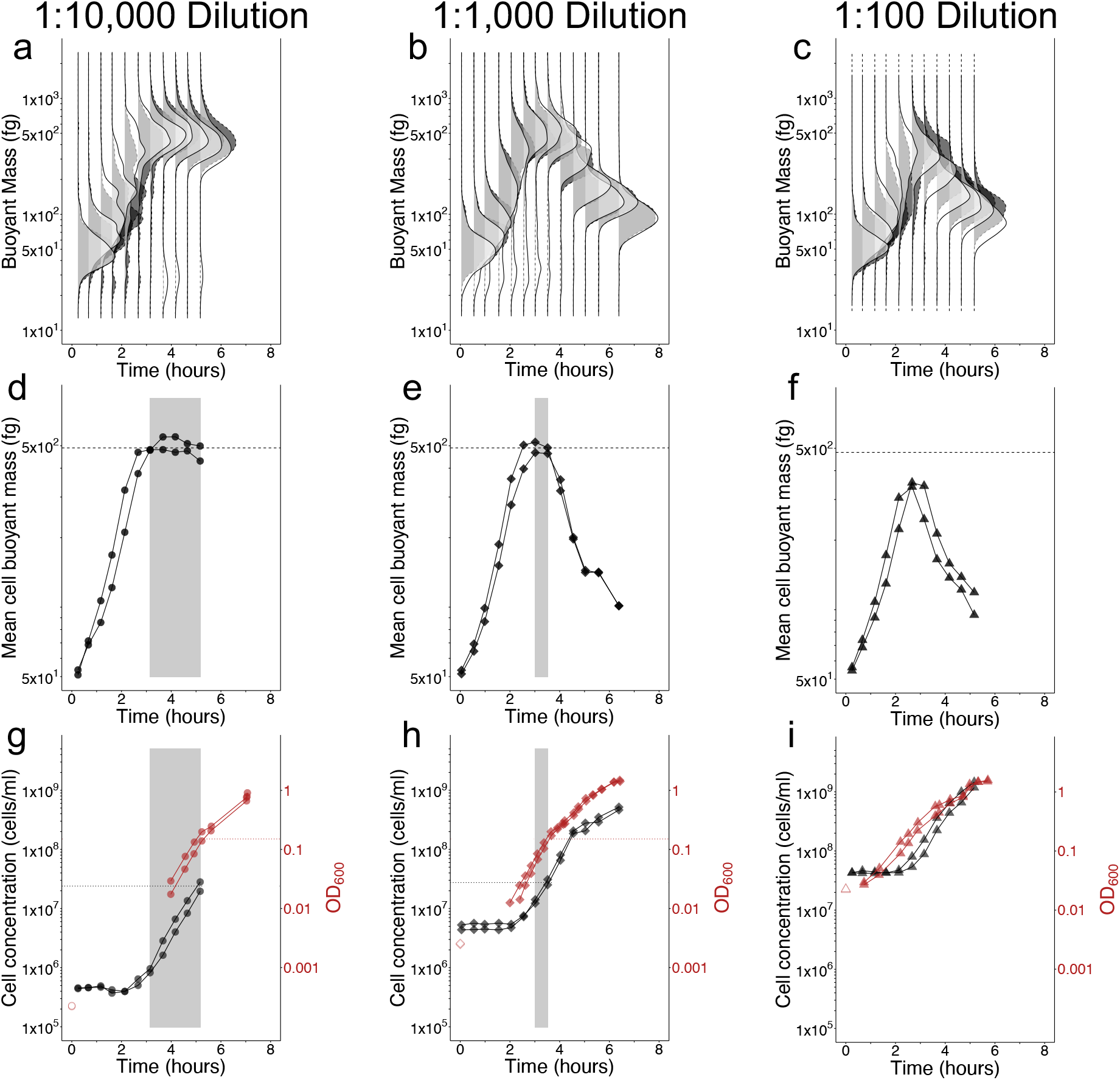
Steady-state growth of *V. cyclitrophicus* in Marine Broth 2216 (MB2216) batch cultures is short-lived and depends on inoculum concentration. Buoyant mass distributions (a-c), mean buoyant mass (d-f), cell concentration (g-i), and optical density 600nm (OD_600_) (g-i) for 2 biological replicates (2 shades of gray a-c, 2 symbols d-i) of cultures inoculated into MB2216 medium with varying dilutions of a stationary phase culture (circles 1:10,000 a,d,g; diamonds 1:1,000 b,e,h; or triangles 1:100 c,f,i). Gray boxes (d, e, g, h) illustrate time periods when steady-state growth was effectively achieved (if ever) based on a constant population mean buoyant mass (dashed line in d, e, f is grand mean of the mean buoyant mass values in the gray boxes). Solid lines drawn between points (d-i) connect consecutive observations for each replicate. Dotted lines indicate the OD_600_ and cell concentration values at the time the culture exits steady-state growth. Unfilled red symbols are provided as reference for the expected inoculum starting OD_600_ based on the measured OD_600_ of the stationary-phase parent culture at the time of inoculation and the known dilution factor, since the 1:1,000 and 1:10,000 dilutions brought the starting OD_600_ of experimental cultures below the lower limit of accuracy for the spectrophotometer (0.01 units OD_600_). Sample size of cells per replicate and timepoint provided in supplemental table 1.

**Figure 2:**
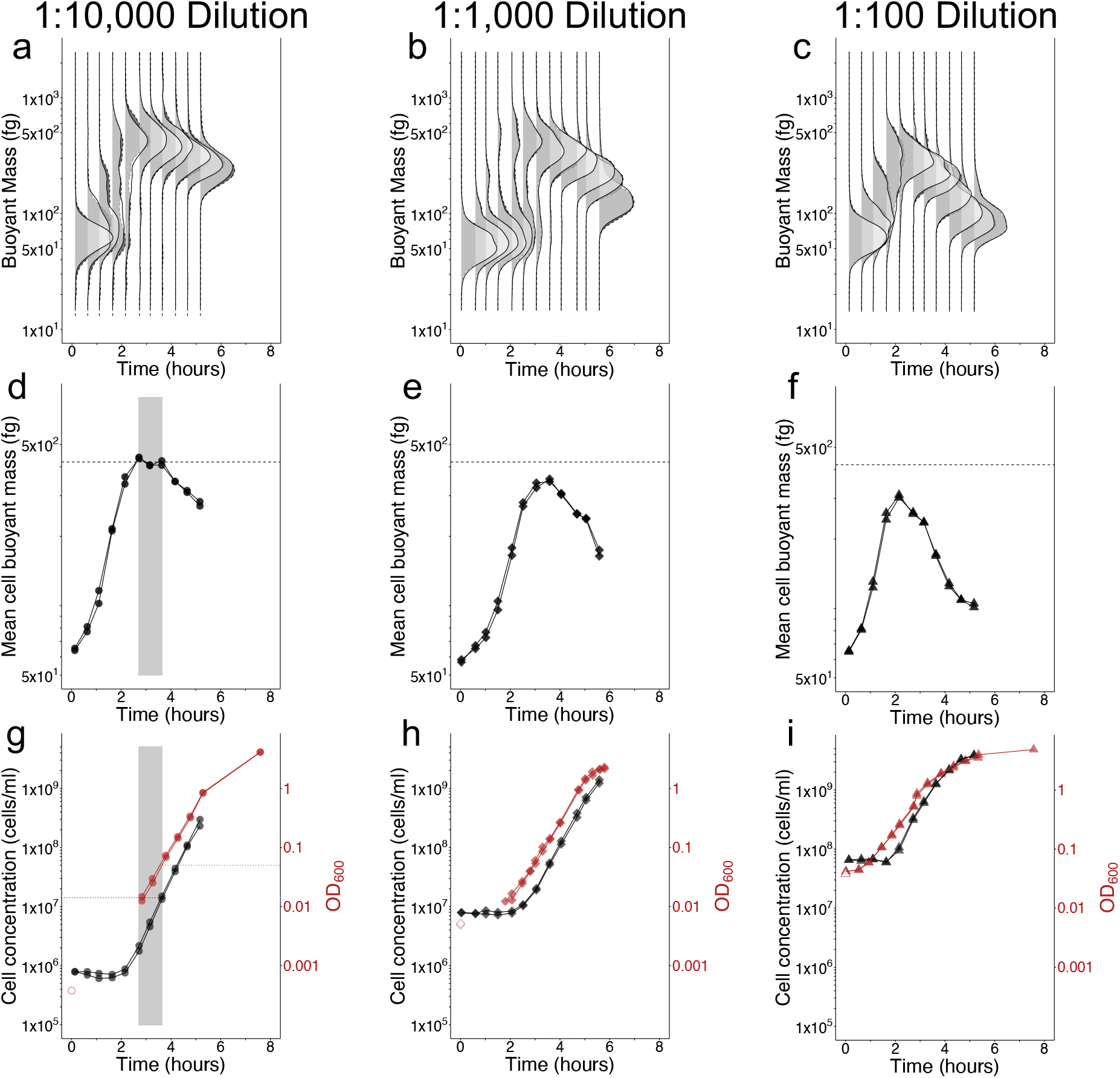
Steady-state growth of *E. coli* in Lysogeny Broth (LB) batch cultures is short-lived and depends on inoculum concentration. Buoyant mass distributions (a-c), average buoyant mass (d-f), cell concentration (g-i), and optical density 600nm (OD_600_) (g-i) for 2 biological replicates (2 shades of gray a-c, 2 symbols d-i) of cultures inoculated into LB medium with varying dilutions of a stationary phase culture (circles 1:10,000 a,d,g; diamonds 1:1,000 b,e,h; or triangles 1:100 c,f,i). Gray boxes (d, g) illustrate time periods when steady-state growth was effectively achieved (if ever) based on a constant population mean buoyant mass (dashed line in d, e, f is grand mean of the mean buoyant mass values in the gray box). Solid lines drawn between points (d-i) connect consecutive observations for each replicate. Dotted lines indicate the OD_600_ and cell concentration values at the time the culture exits steady-state growth. Filled symbols represent measured data. Unfilled red symbols are provided as reference for the expected inoculum starting OD_600_ based on the measured OD_600_ of the stationary-phase parent culture at the time of inoculation and the known dilution factor, since the 1:1,000 and 1:10,000 dilutions brought the starting OD_600_ of experimental cultures below the lower limit of accuracy for the spectrophotometer (0.01 units OD_600_). Sample size of cells per replicate and timepoint provided in supplemental table 1.

The number of OD, or total mass, doublings required for reaching steady-state was fewer than we assumed based on typical literature guidelines of 10 doublings(2, 16). At the time steady-state begins, the average cell had undergone about 3 mass doublings (*V. cyclitrophicus:* 53 to 492fg, *E. coli*: 63 to 420fg), while the cell concentration had doubled (Figure 1 g & h, Figure 2g) combining to 4 total mass doublings. This agrees well with the increase in OD from inoculation until steady-state onset (Figure 1h, Figure 2g). We have shown the number of doublings necessary for ensuring steady-state growth can be as low as 4 when diluting a stationary phase culture into the same medium. Other single-cell studies examining different transitions between steady-states found it takes 3-8 doublings to establish the new steady-state after a growth medium shift(19, 20), but they do not try to examine if their findings can be translated to traditional batch culture and optical density growth. Taken together this suggests the guideline of 10 doublings for ensuring steady-state establishment is robust, but may be overly cautious.

The amount of mass gained by cells is enormous as they transition into steady-state growth following starvation. While the mean buoyant mass change we observed could be influenced by changes to the composition of cellular dry mass via altered dry density, this effect is relatively small for *E. coli* in similar conditions(17). Therefore, *E. coli* buoyant mass can be converted to dry mass using a multiplicative conversion factor between 3.1 (in stationary phase) to 3.6 (near steady-state)(17). *E. coli* alters its dry mass more in the first 3 hours of our experiments (164fg-1,512fg), than it does across 20 different growth media supporting steady-state growth rates between 0.31-1.72 per hour (240fg-1,180fg)(16). Experimentalists comparing mass in multiple conditions (e.g., different media) must account for this within condition variation by establishing steady-state growth. The mass range clonal isolates can realize is vast, but cell mass is systematically responsive to the transient and long-term(16) nutrient conditions bacteria experience.

The exit from steady-state occurs very early in the growth curve at low OD and cell concentration. This has been reported previously for *E. coli* in LB(4, 5) and we observe steady-state exit to occur at an even lower OD and cell concentration (Figure 2g: ∼0.1 units OD_600_ and ∼1.5×10^7^ cells/ml). This early exit phenomenon seems to be generally true in rich medium, as *V. cyclitrophicus* also exits at low OD and cell concentration in both dilution treatments (Figure 1 g & h: ∼0.1 units OD_600_ and 2.5×10^7^ cells/ml). Both species OD growth curves have subtle bends just after steady-state departure (Figure 1g & h, Figure 2g), which has been interpreted for *E. coli* in LB as the start of a sequential depletion of easily catabolized amino acid equivalents(5). While the absolute exit values for OD and cell concentration are similar to one another for the two species, the final yield of these species in their respective media differs (Final yield: *E. coli* = ∼5 units OD_600_ and ∼7-8×10^9^ cells/ml; *V. cyclitrophicus* = ∼2 units OD_600_ and ∼5×10^9^ cells/ml). We also note that steady-state growth is only achieved if the concentration of cells at inoculation is at least 1 doubling below the cell concentration at which the population exits steady-state.

Although population averages are of central importance to steady-state phenomena, the mass distribution itself and how it shifts in these experiments can provide additional insights into cell individuality in fluctuating conditions. Mass initially appears to have a log-normal distribution upon inoculation into fresh medium and it quickly broadens (increased robust coefficient of variation, Supplemental Figure 1 a-c, Supplemental Figure 2 a-c,) while the central tendency rapidly shifts upwards (Figure 1 a-c & Figure 2 a-c). This indicates that some cells add mass more quickly than others. There is also no obvious sub-population of non-responsive cells since the mass distribution at its maximum (2-3 hours) is entirely larger than the initial mass distribution (Figure 1 a-c & Figure 2 a), though they may be too rare to detect with the number of cells we observed. The mass distribution upon entry into steady-state returns to a more narrow log-normal distribution (Figure 1 a-c & Figure 2 a) with a smaller rCV (Supplemental Figure 1 a-c, Supplemental Figure 2 a-c), which is the expected distribution for cultures growing in steady-state growth(21). Upon exiting steady-state, the center of the distribution decreases but does not broaden (Figure 1 a-c & Figure 2 a-c) or have a change in rCV (Supplemental Figure 1 a-c, Supplemental Figure 2 a-c) indicating no change in variation among cells as they reductively divide to make offspring with ever decreasing size.

## Conclusions

Microbiologists have known for a century that cell properties in liquid batch cultures are dynamic(6), but exactly how much cells alter their mass over time is challenging to measure directly. We believe this information gap, along with misconceptions about OD have led to the common misunderstanding that an exponential OD increase is equivalent to steady-state growth. Here we demonstrate that cell mass changes rapidly and substantially in batch culture, so care must be taken to achieve steady-state growth. While the exact amount of mass added upon dilution into fresh medium will depend on the specific strain and medium combination, it is a very large increase for these strains in commonly used rich medium.

**We recommend that microbiologists intending to work with steady-state populations in batch culture on rich media perform the following steps:**

**1) inoculate with a minimum of a 10**,**000-fold dilution from an overnight parent culture for *E. coli*-like cell yields (∼10**^**9**^ **cells/ml)**.

**2) allow for at least 4 OD, or total mass, doublings (16-fold) prior to initiating an experiment**.

**3) finish the experiment at 0.1 units OD**_**600**_ **& before the earliest bend in the growth curve**.

One major caveat is that OD measurements are never exactly comparable across devices or with different growth media & strain combinations(9). We believe our guidelines are cautious minimum standards and illustrate the complex interplay between individual and population growth all bacteria face in batch culture. We recognize that these criteria require an experimental decision about observing OD changes. This decision arises because the lower accuracy limit of OD in many spectrophotometers is near 0.01(22), and starting from this point does not allow for 4 mass doublings, and therefore entry to steady state, before the OD threshold of 0.1 is reached. One can either start from the OD of 0.01 and perform repeated dilutions of the culture until reaching 4 mass doublings, or start from outside of your observation window by performing a larger dilution. When performing a larger dilution, using the known OD of the parent culture and the dilution factor can reliably measure starting OD and ensure at least 4 total mass doublings finish in the range of 0.01-0.1 OD.

The mass dynamics we documented impact all experiments using batch cultivation, from comparing gene or protein expression to interpreting the dynamics of ecology and evolution experiments. For example, a common experimental design for studying microbial evolution is the serial daily passage of a batch culture with a 1:100 dilution(23). These experiments frequently observe strong selection on cell size and growth rate, likely driven by a rapid mass increase upon inoculation into fresh medium similar to our findings(24, 25). Those cells which increase in mass fastest will have a large advantage compared to slow responding cells. Serial passage experimental designs have also been used to study ecological dynamics within multispecies microbial communities(26, 27). All serial passage experiments likely face similar selective pressures on cell size and growth rate, while individual species may exhibit different mass accumulation responses. Finally, based on final yield and our measurements we see that the majority of total cell divisions in all of our dilution cultures occur outside of steady-state. Therefore, the dynamics observed between co-occurring cells in batch culture experiments will be strongly influenced by the physiology of mass accumulation and contrasting facets of individual vs. population growth.

## Methods

### Bacterial strains and culture conditions

*Escherichia coli* K12 MG1655 containing the pSIM6 plasmid(28) was provided by the laboratory of Dr. Robert Britton and grown in autoclaved & 0.1µm filtered Lysogeny broth (LB Lennox) liquid medium (Carl Roth GmbH). *Vibrio cyclitrophicus* 1G07 was isolated from cryopreserved seawater samples as previously described(29) and grown in boiled and 0.1µm filtered Marine Broth 2216 (BD Difco) liquid medium (Fisher Scientific GmbH). *E. coli* was grown in pre-warmed LB at 37°C shaking at 200 RPM while *V. cyclitrophicus* was grown at 25°C shaking at 250 RPM. The following cultivation protocol was followed for each experiment. For each species, two biological replicate cultures were inoculated on day 1 from the same -80°C freezer stock into separate recovery cultures (5ml medium in 13ml test tube) and allowed to grow overnight. We define biological replicate as an independent culture originating from the same clonal freezer stock, but which was inoculated into a separate recovery culture and maintained separately for all further transfers. On the morning of day 2 the recovery cultures had all achieved stationary phase and high optical density, so were inoculated with a 1:100 dilution into pre-cultures (50µl into 5ml medium in 13ml test tube) and allowed to grow for an additional day. This ensured the cultures had several generations of growth to recover from cryopreservation and were also in stationary phase in the same medium used for experimentation for about 18 hours. Experimental cultures were inoculated on day 3 using different dilutions (either 1:100, 1:1,000, or 1:10,000) of stationary phase pre-cultures into 30ml of pre-warmed medium. Experiments with the 1:100 and 1:10,000 dilutions were performed on the same day and with the same 2 biological replicate pre-cultures as inoculant sources. These experimental cultures were inoculated serially, first diluting the pre-cultures to generate the 1:100 experimental cultures (0.3ml pre-culture into 29.7ml medium in 500ml side-arm flasks) and then further to generate the 1:10,000 experimental cultures (0.3ml of 1:100 culture into 29.7ml medium in 500ml side-arm flasks). Experiments with the 1:1,000 experimental cultures were performed several months later with the same procedure until the inoculation of experimental cultures, where the replicate pre-cultures were diluted to generate the 1:1,000 experimental cultures (30µl pre-culture into 30ml medium in 500ml side-arm flasks).

### Optical density measurements

Optical density at 600nm (OD_600_) was measured on a Genesys40 spectrophotometer (Thermo Fisher) relative to uninoculated growth medium blanks in either cuvettes or test tubes where appropriate. The accuracy limits of the spectrophotometer (0.01-0.79 OD_600_) were determined with a dilution series of a stationary phase culture of *V. cyclitrophicus* 1G07 which was washed and re-suspended in a buffer that did not contain the macronutrients necessary for growth. OD_600_ of cultures was measured in the side-arm of the cultivation flask until approaching an OD_600_ value around 0.5, after which an additional 100µl of sample was destructively removed from the flask to dilute and measure the true value withing the accuracy limit of the spectrophotometer. Briefly, these samples were diluted 1:10 with fresh medium (0.1ml sample into 0.9ml medium) in a cuvette and immediately measured in the same spectrophotometer with a cuvette adapter. Diluted sample OD_600_ values were then multiplied by the dilution factor.

### Single-cell mass and cell concentration measurements

Single-cell mass measurements were made on the LifeScale-Research instrument (Affinity Biosensors). The sensor on this device is a suspended microchannel resonator similar to previously described and published devices(30). The device was configured with automated sampling hardware and software and its flow speed optimized for resolving particles at the smallest size possible (around 14fg buoyant mass). Calibration curves with NIST certified polystyrene beads of known diameter and density were used to verify the accuracy and precision of mass measurements.

Batch cultures were prepared with medium that was filtered with 0.1µm pore size PES filters immediately prior to use to ensure it was free of any small particles that could interfere with cell measurements on the SMR cantilever. Negative controls of uninoculated medium were run on the SMR prior to inoculation to ensure the particle background of each medium batch was adequately low (concentrations of less than 1×10^5^ background particles/ml, comparable to ultrapure water controls).

Samples were removed from batch cultures immediately after inoculation, and at approximate 30-minute intervals for approximately 5-6 hours. For 1:10,000 and 1:100 dilution cultures 11 samples were taken over 5 hours. For 1:1000 dilution cultures 13 samples were taken over 6 hours. Sample fixation was performed by removing 1ml of culture and adding it to pre-aliquoted microcentrifuge tubes containing 0.205ml of ice-cold, 0.1µm PES filtered formaldehyde (4% final concentration, from 23.5% methanol-free formaldehyde solution). Fixation samples were kept on ice for 1 hour before storing permanently at 4°C. The measurement of samples for single-cell mass and cell concentration was performed at least 1 day after samples were fixed. The concentration of cells in successive time point samples changed over time due to growth, but the LifeScale sensor can only operate accurately at concentrations between approximately 1×10^4^-3×10^7^ cells/ml. Therefore, we removed a small volume from each fixation sample after vortexing to resuspend cells, and diluted it in 0.1um PES filtered growth medium (where necessary) to target a concentration of 1×10^7^ cells/ml. Measured concentrations were then adjusted for this sample dilution. We do note the 1:10,000 cultures were below the 1×10^7^ cells/ml target concentration until near the end of the experiment, so many samples from these cultures were counted without any dilution. This resulted in fewer cells counted per sample of the 1:10,000 cultures than the 1:100 cultures or the 1:1,000 cultures. Sample sizes of cells per sample for figures 1 and 2 are included in supplemental table 1. Sample sizes differ slightly for mass (Figures 1 & 2 a-f) and concentration (Figures 1 & 2 g-i) due to some particles passing through the mass sensor nearly simultaneously. If 2 particles pass through the sensor simultaneously and only one particle’s mass can be accurately measured, the software registers 1 particle for mass purposes (1 measured particle) but 2 particles for concentration purposes (2 detected particles). Therefore, the discrepancy in sample size for different figure panels measured from the same sample relates to the difference between measured and detected particles.

### Data analysis and statistics

LifeScale software automatically converts the shift in cantilever oscillation frequency into buoyant mass values for each cell. It also detects and reports the fluid flow rate and measures the exact fluid volume and particle number to calculate cell concentration in a sample. Cell concentration and buoyant mass data was exported from LifeScale software in .csv format. All further analysis, statistics, and figure generation was performed in the R statistical programming language using base R(31), the tidyr package(32), and ggridges(33).

## Data availability statement

Data used in this study and code necessary to re-generate all figures will be available publicly on the Polz Lab github (https://github.com/polzlab) upon publication of this manuscript.

## Acknowledgements

Former Polz lab members Kathryn Kauffman, Joseph Elsherbini, and Fatima Hussain for collecting seawater time series samples and isolating the *V. cyclitrophicus* 1G07 strain used in this study. S.R.M. is a co-founder of Travera and Affinity Biosensors, which develop technologies relevant to the research presented in this work. This work was supported by the Simons Foundation (Life Sciences Project Award – 572792) to M.F.P..

